# fgf8a signalling shapes brain divergence between Malawi cichlids

**DOI:** 10.1101/2025.10.13.682009

**Authors:** Aleksandra Marconi, Jake Morris, Pío Sierra, Dillan Saunders, Joel Elkin, Benjamin Steventon, Richard Durbin, Stephen H. Montgomery, M. Emília Santos

## Abstract

Brain diversification is intimately linked with adaptive radiations, yet the underlying molecular basis remains poorly understood. Here, we examine the mechanisms of neural evolution in two ecologically divergent Lake Malawi cichlid species: a generalist, *Astatotilapia calliptera*, and a pelagic piscivore, *Rhamphochromis* sp. ‘chilingali’. We demonstrate that forebrain domains diverged independently from other brain regions in these species, consistent with mosaic brain evolution. We identify *fibroblast growth factor 8a (fgf8a)* as a key factor underlying this neuroanatomical divergence. Species-specific *fgf8a* expression patterns during critical developmental windows correlate with adult brain differences. Functional knockout experiments confirm *fgf8a*’s essential role in cichlid brain patterning, directly linking this conserved developmental gene to interspecific neuroanatomical variation. We identify species-specific transposable element repertoires in the Malawi cichlid *fgf8a* locus that function as developmental enhancers in brain and sensory systems, providing a potential mechanism for expression divergence between species. Phylogenetic analysis across the radiation reveals elevated transposable element polymorphism at the *fgf8a* locus in *Rhamphochromis*, suggesting ongoing regulatory evolution in this clade. Our findings reveal how conserved developmental programmes can accommodate regulatory modification, providing a mechanistic framework for understanding rapid neural diversification during adaptive radiation.

## Introduction

Brains and sensory systems exhibit remarkable diversity across vertebrates, reflecting their pivotal roles in mediating species-specific ecological and behavioural adaptations (1). Among teleosts, which comprise ∼96% of extant fish species (2), African Great Lakes cichlids stand out as a powerful model for exploring mechanisms underlying rapid neural diversification. Their iconic, large-scale adaptive radiations in Lake Malawi, Tanganyika and Victoria, each encompass hundreds of closely related species that differ in habitat use, trophic ecology, and behavioural strategies (3,4). Importantly, Eastern African cichlid radiations showcase neuroanatomical variation that is closely associated with ecological niche and sensory modality (5–7), making them an ideal system for dissecting the genetic and developmental underpinnings of adaptive nervous system evolution.

Previous studies demonstrated that interspecific differences in cichlid brain morphology are established early in neural development (8–10). Discrete spatio-temporal alterations in conserved signalling pathway expression are associated with variation in the relative size of forebrain regions between sand-dwelling (‘nonmbuna’) and rock-dwelling (‘mbuna’) species (8,10). However, the developmental, genetic and regulatory mechanisms underlying ecologically relevant variation in brain morphologies remain poorly understood.

In this study, we focus on *fibroblast growth factor 8a* (*fgf8a*), a morphogen with essential roles in vertebrate brain and sensory development (11–14). This candidate initially emerged from previous transcriptomic and population genetics analyses in two divergent species from Lake Malawi basin: the shallow weedy-water generalist, *Astatotilapia calliptera*, and open-water pelagic piscivore, *Rhamphochromis* sp. ‘chilingali’ (15) (Fig. 1A-B). Two sources of evidence implicate *cis*-regulatory elements at *fgf8a* in adaptive divergence between these species. First, *fgf8a* was differentially expressed between species at multiple stages during embryogenesis. Second, a genome-wide selection scan identified a strong cross-population extended haplotype homozygosity (xp-EHH) signal in the upstream non-coding region of *fgf8a* (Fig. 1C). In zebrafish and other vertebrates, *fgf8* signalling is required for brain regionalisation at multiple organiser centres located at the borders between forebrain, midbrain and hindbrain (16–17). Subtle variation in embryonic expression of *fgf8a* has previously been identified between sighted and blind morphs of cavefish (*Astyanax mexicanus)* (18), but their phenotypic relevance is unclear. Beyond functions in brain regionalisation, fgf8 signalling is also involved in development of several sensory traits in teleosts, particularly visual and auditory systems (19–22). Finally, the complexity of *fgf8* regulation, comprising dozens of *cis*-regulatory elements distributed over extensive genomic range, has been reported across vertebrates (23). These elements are involved in precise localisation and dosage as well as context-dependent modulation of *fgf8a* activity (2, 7, 14, 23, 24).

**Figure 1.**
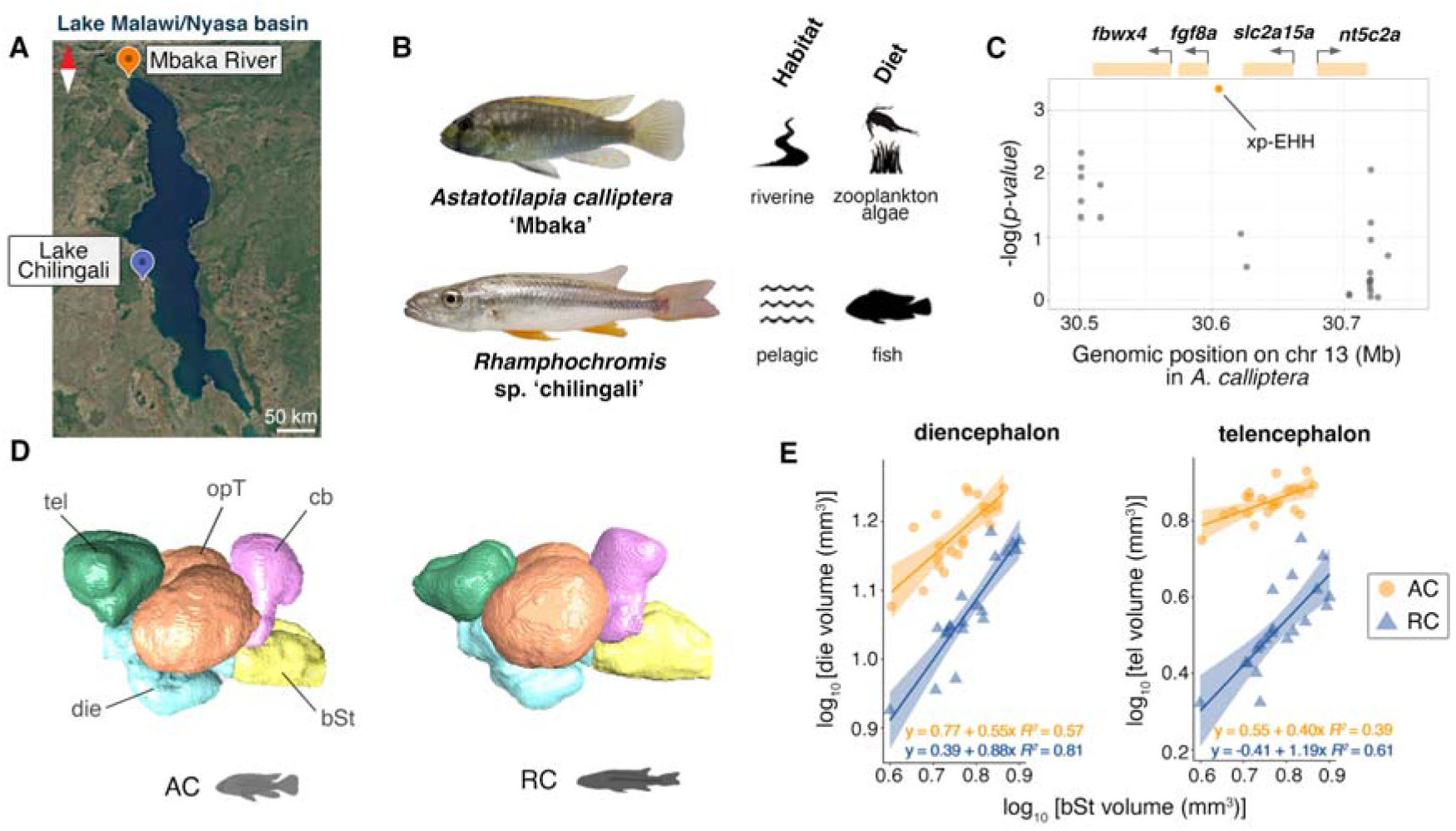
Ecological, genomic, and neuroanatomical divergence correlates in cichlids of Lake Malawi basin. **A.** Map of Lake Malawi/Nyasa basin, including Lake Chilingali and River Mbaka. **B.** Lake Malawi basin cichlid species *Astatotilapia calliptera* ‘Mbaka’ (AC) and *Rhamphochromis* sp. ‘chilingali’ (RC) part of this study are characterised by distinct habitat (riverine vs. pelagic) and diets (zooplankton and algae vs. fish). **C.** Our previous work (15) identified a significant recent positive selection signal (cross-population extended haplotype homozygosity, xp-EHH) in non-coding region on chromosome 13, pinpointing *fgf8a*, a key brain patterning morphogen, as potentially involved in adaptive neuroanatomical divergence between AC and RC. The same study identified *fgf8a* as differentially expressed between these species at multiple timepoints during embryonic development. **D.** Three-dimensional volumetric models of representative AC and RC adult brains reconstructed from μCT scans showing segmented subdivisions in lateral views: telencephala (tel), optic tecta (opT), diencephala (die), rhombencephala (including dorsally protruding cerebellum, cb) and brain stems (bSt). **E.** Significant variation in the diencephalon and telencephalon volumes between AC (circles) and RC (triangles). Scaled (log-transformed) proportions against the volume of the brain stem to control for brain size differences with linear regressions indicating species-specific neuroanatomical scaling relationships. Shaded bands correspond to 95% confidence intervals. N > 15 for both species. Map in (**A**) after Google Earth (2025). Photographs in (**B**) modified from ref. 15 with authors’ permission. Abbreviations: AC - *Astatotilapia calliptera* ‘Mbaka’, bSt - brain stem, cb - cerebellum, die - diencephalon, opT - optic tecta, RC - *Rhamphochromis* sp. ‘chilingali’, tel - telencephalon, xp-EHH - cross-population extended haplotype homozygosity.

While a causal role for *fgf8* in the evolution of forebrain architectures has not been explicitly demonstrated in any species to date, we hypothesised that subtle differences in *fgf8a* expression and regulation could therefore have disproportionate effects on the regional boundaries between brain regions. To test this hypothesis, we performed anatomical, developmental and genetic comparisons between *Astatotilapia calliptera* and *Rhamphochromis* sp. ‘chilingali’. We first demonstrate that the adult differences between focal species involve volumetric differences in forebrain territories (diencephalon and telencephalon) canonically patterned by *fgf8a.* We then explore the role of *fgf8a* in cichlid neural divergence. We show that *fgf8a* expression patterns during embryonic brain morphogenesis differ markedly between examined species, in both localisation and spatial domain sizes. CRISPR/Cas9-mediated disruption of *fgf8a* in *A. calliptera* produces broad neuroanatomical defects indicating functional relevance in brain development. Comparative genomic analysis further revealed multiple transposable element (TE) insertions specific to the Malawi radiation and *A. calliptera*, and we demonstrate that LINE-2 and PIF-Harbinger insertions can function as developmental enhancers with distinct species-specific activity patterns in neural and sensory tissues. By integrating genomic, developmental, and evolutionary perspectives, our study provides new insights into the molecular underpinnings of nervous system diversity in one of nature’s most iconic adaptive radiations.

## Results

### Variation in brain morphology between ecologically divergent Malawi cichlids

Brain morphology is expected to reflect adaptations to divergent selective pressures imposed by distinct environments. To investigate this phenomenon in cichlids, we focused on species belonging to two divergent ecotypes: *Astatotilapia calliptera* ‘Mbaka’ (shallow water generalist, AC) and *Rhamphochromis* sp. ‘chilingali’ (pelagic piscivore, RC) (Fig. 1A-B). *Astatotilapia* is also considered to closely resemble the ancestor of the Malawi radiation (25). Using μCT scans of adult specimens, our volumetric analyses revealed significant differences in the relative size of two forebrain (prosencephalon) regions: the diencephalon and telencephalon (Fig. 1D, Supplementary Fig. 1). Both structures show non-allometric expansion in *A. calliptera* compared to *Rhamphochromis*, demonstrating specific shifts in investment that are independent of overall brain size (diencephalon F(1, 37) = 4.820, *p* = 0.034; telencephalon F(1, 37) = 9.366, *p* = 0.004, one-way analysis of variance, Fig. 1E), and not observed for other structures (Supplementary Fig. 1). This adult variation supports a model of mosaic brain evolution, where individual brain regions evolve independently rather than as a single, coordinated unit (26), in ecological diversification in cichlids (6, 27–29). Our data further suggest that the morphogenesis of the diencephalon and telencephalon diverge between *A. calliptera* and *Rhamphochromis* during development and underscore the need for further investigation into brain development during embryogenesis to elucidate the underpinnings of this adaptive variation.

### Quantitative variation in *fgf8a* expression during brain development between cichlid species

The signatures of positive selection we identified in *fgf8a*’s regulatory region suggest that expression divergence during development may underlie adult brain differences between species. We therefore used Hybridization Chain Reaction (HCR) *in situ* hybridization (30) to quantify *fgf8a* expression throughout embryonic development, focusing on critical stages when brain regional identity is established.

Comparative analysis revealed dynamic species-specific expression differences that emerge early in development and persist throughout brain regionalisation. During formation of early brain rudiment stage (4 somite stage, ss), the species exhibit divergent *fgf8a* expression patterns: *Rhamphochromis* shows restricted hindbrain expression, while *A. calliptera* displays broader anterior neural tube distribution, including the presumptive forebrain domain (Fig. 2B). This early divergence suggests species-specific variation in anterior neural patterning and cranial placode specification.

**Figure 2.**
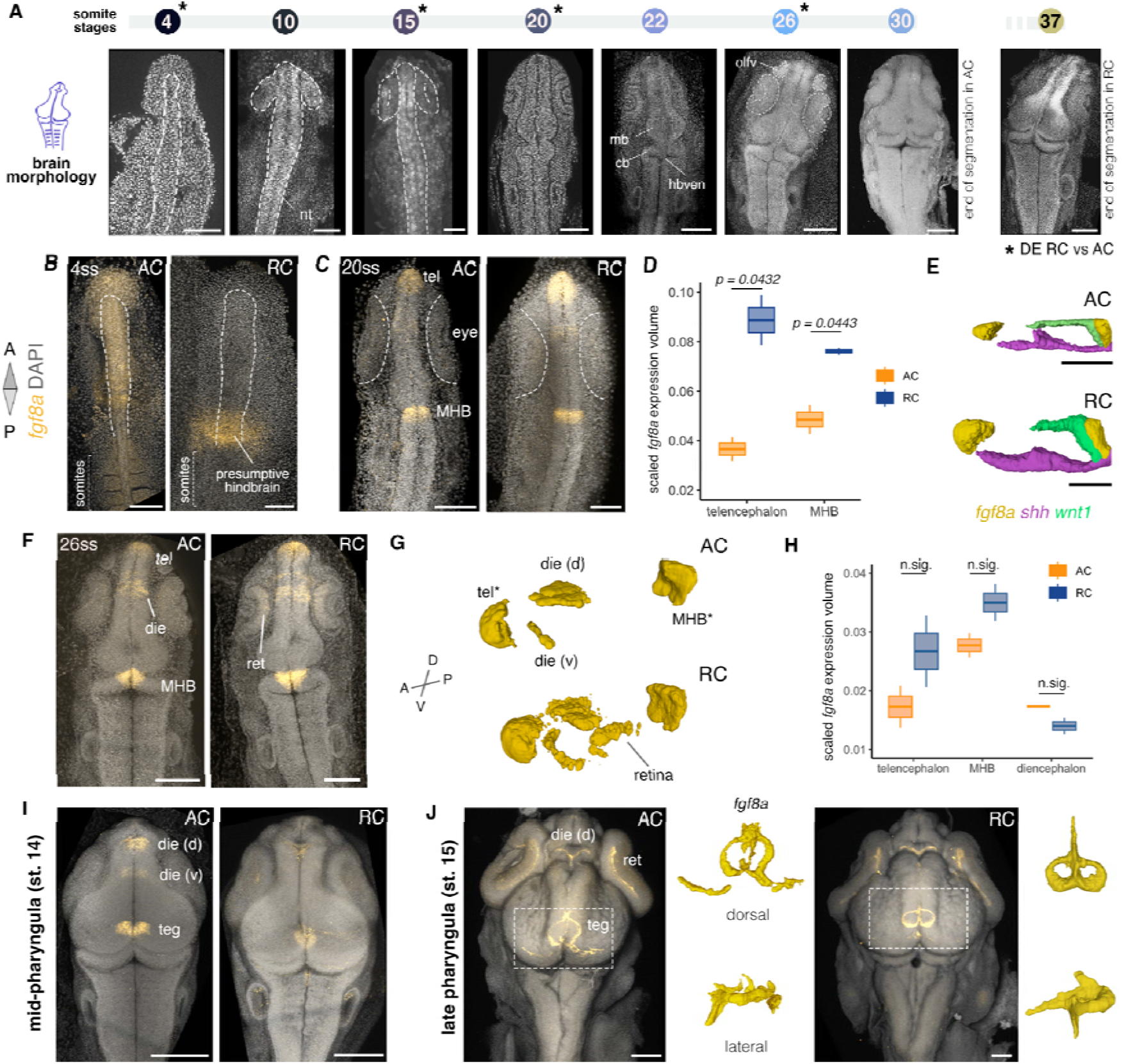
Quantitative variation in *fgf8a* expression during embryonic brain development between cichlid species. **A.** Overview of early brain morphogenesis in *Astatotilapia calliptera*, ranging from early neurulation (4ss) until regional differentiation of the brain (end of segmentation), concluding at 30 and 37ss in *A. calliptera* and *Rhamphochromis*, respectively. Our previous analysis detected differential expression of *fgf8a* between study species across multiple developmental stages (15), indicated here with asterisks. Dorsal views of embryos dissected from yolk and stained with nuclear marker DAPI. Anterior to the top. **B.** Early divergence in anterior neural tube patterning by *fgf8a* between species at 4ss. **C-E.** Differences in *fgf8a* expression and brain patterning are evident at 20ss. **C.** *fgf8a* is expressed in telencephalon and MHB in both species. **D.** These two *fgf8a* domains do not scale proportionally with total brain volume in these species at 20ss (Student’s t-test, MHB t(2) = -4.59, *p* = 0.0443, telencephalon t(2) = -4.65, *p* = 0.0432). Independent biological replicates *n* = 3 per species. **E.** 3D renditions of expression domains of signalling molecules *wnt1* and *shh* reveal variation across multiple developmental programmes. Lateral views. **F.** At 27ss, *fgf8a* expression is now also detected in dorsal and ventral diencephalon in both species as well as in retina in RC. **G.** 3D volume renditions of *fgf8a* expression domains at 27ss.**H.** Scaling differences involving *fgf8a* territories between species do not persist into late somitogenesis (Student’s t-test, MHB *p* = 0.193, telencephalon *p* = 0.314). Independent biological replicates *n* = 3 per species. **I-J.** *fgf8a* expression during pharyngula stages includes distinct expression patterns in the tegmentum. 3D volume renditions of tegmental *fgf8a* expression (in dashed boxes) in dorsal (top) and lateral (bottom) views presented in side panels in J. All images show representative examples of embryos from at least three replicates. Abbreviations: AC - *Astatotilapia calliptera*; d - dorsal; DE - differential expression; die - diencephalon; MHB - midbrain-hindbrain boundary; RC - *Rhamphochromis* sp. ‘chilingali’; ss - somite stage; st - stage; teg - tegmentum; tel - telencephalon; v - ventral. All scale bars equal to 100μm.

By 20ss, when major brain compartments are clearly demarcated, *fgf8a* expression was established in canonical signalling centres, including the midbrain-hindbrain boundary (MHB) and telencephalic anterior neural ridge (ANR) in both species (Fig. 2C). However, quantitative analysis revealed that *Rhamphochromis* exhibited larger *fgf8a* expression domains in both regions compared to *A. calliptera* at 20ss (Student t-test, p < 0.05) (Fig. 2D). These differences did not persist through late somitogenesis (27ss), albeit *Rhamphochromis* showed retinal expression absent in *A. calliptera* at this stage (Fig. 2F-H). The observed volumetric differences during early brain development point towards changes in the spatial distribution of territories influenced by *fgf8a* signalling.

Three-dimensional reconstruction analysis revealed that species-specific differences extended beyond *fgf8a* to encompass other key signalling molecules within the developing brain (Fig. 2E). The posteriorizing signal *wnt1* showed reduced midbrain expression in *Rhamphochromis*, while the zona limitans intrathalamica (ZLI) marker *shh* exhibited greater dorsal extension at a more anterior position (relative to midbrain-hindbrain boundary) in *Rhamphochromis*. These expression patterns indicate coordinated species-specific modifications to the molecular networks establishing forebrain-midbrain-hindbrain boundaries, with AC showing enlarged posterior diencephalic territory relative to *Rhamphochromis*.

During pharyngula stages, *fgf8a* expression becomes increasingly regionalised, with the most pronounced species differences emerging in the developing tegmentum. At mid-pharyngula stage (st. 14, Fig. 2I) AC exhibited stronger signals in the dorsal diencephalon (presumptive pineal complex), whereas by late pharyngula (st. 15, Fig. 2J), AC showed an expanded, irregular *fgf8a* domain with bilateral lateral extensions in AC compared to *Rhamphochromis*’ compact, bilaterally symmetric pattern. These late-appearing differences may contribute to species-specific neural circuit organisation and sensory system development influenced by fgf8a signalling.

The spatio-temporal expression analyses reveal that *fgf8a* exhibits dynamic, species-specific deployment throughout cichlid brain development. While conserved expression in canonical signalling centres (MHB, ANR) underscore the conserved role of fgf8(a) signalling in vertebrate brain regionalisation, the pronounced interspecific variation in expression levels and spatial distribution suggests cumulative modifications to *fgf8a* developmental programmes. As fgf8a functions as a diffusible morphogen, these expression differences likely generate distinct concentrations gradients that pattern brain territories beyond immediate expression domains, as well as influence the interactions with other signalling pathways, potentially explaining how early molecular divergence translates into the adult volumetric differences observed between species. Given that morphogen effects do not necessarily correspond directly to expression domains, functional validation through targeted gene disruption was deemed essential to establish causal relationships between *fgf8a* activity and the specific brain regions (i.e. diencephalon and telencephalon) showing natural morphological variation between study species.

### Functional disruption of *fgf8a* alters brain, craniofacial and body axis development in cichlids

To directly test whether *fgf8a* differentially affects the development of brain morphology between cichlid species, we employed CRISPR/Cas9 gene editing in *A. calliptera*. Targeted frame-shift mutations in exon 2 (Fig. 3A) created mosaic mutants that allowed us to assess *fgf8a*’s role in cichlid brain development while avoiding early lethality (Supplementary Fig. 2, Supplementary Table 1).

**Figure 3.**
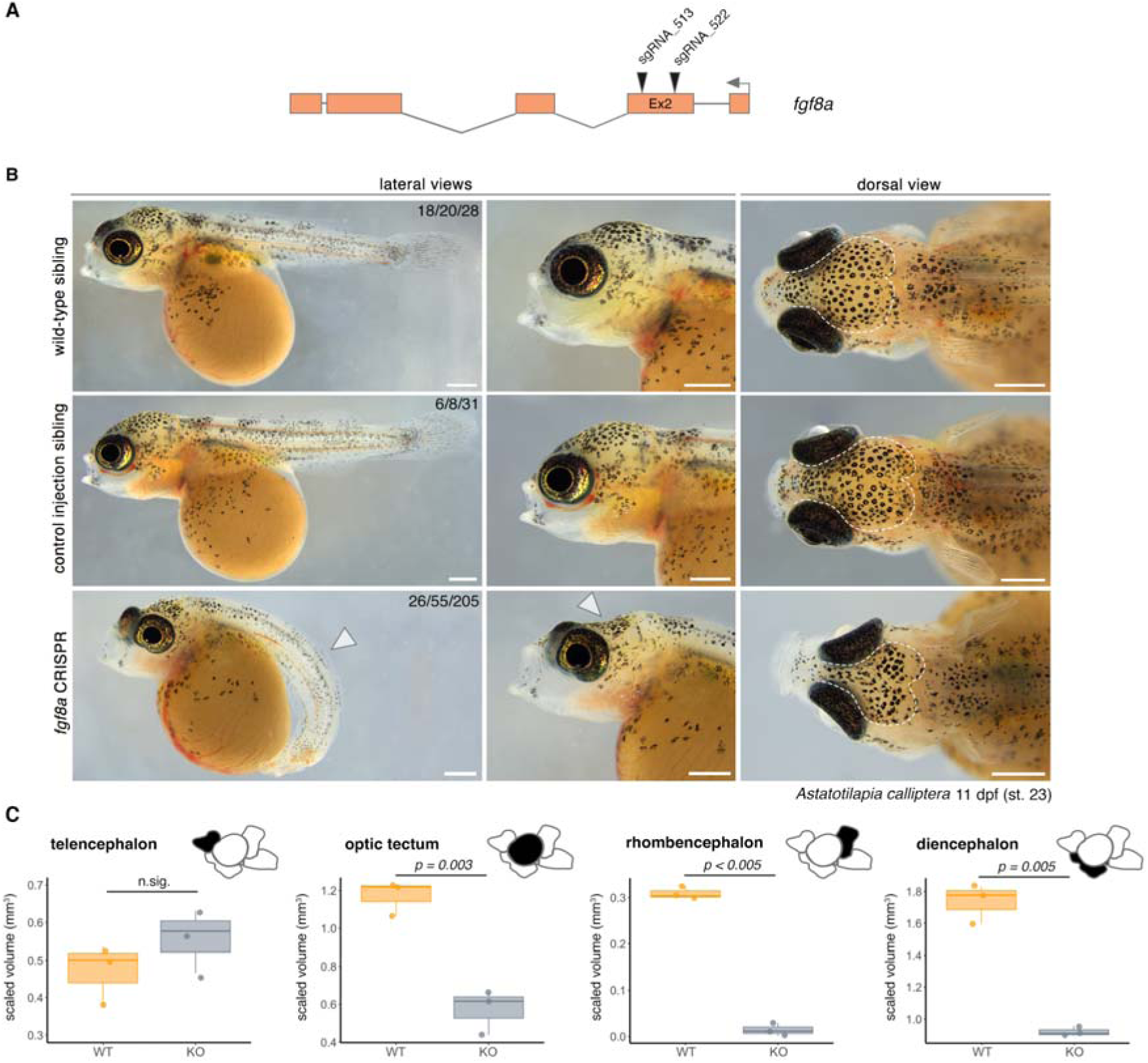
Functional impact of *fgf8a* knockout on cichlid development. **A.** Overview of the *fgf8a* CRISPR/Cas9-mediated knockdown. **B.** Craniofacial and body axis defects in *fgf8a* CRISPR/Cas9 mosaic mutants of *Astatotilapia calliptera*. Representative images of wild-type siblings (top row), control-injected siblings (middle row), and *fgf8a* CRISPR mutants (bottom row) at 11 dpf (stage 23). Lateral views (left) show impaired tail straightening and reduced axial elongation in mutants (white arrowheads). Dorsal views (right) highlight abnormal eye positioning and craniofacial flattening (white arrowheads). Proportions shown indicate phenotype presented/all alive and scored on the day/all injected. **C.** *fgf8a* CRISPR mosaic mutants display significant volumetric reductions in forebrain (diencephalon), midbrain (optic tectum) and hindbrain (rhombencephalon) structures. Scaled proportions against the volume of the brain stem shown to control for brain size differences. Welch Student’s t-test used to test for significant differences in means: telencephalon *p* = 0.275, optic tectum t(3.78) = -6.8773, *p* = 0.0029, rhombencephalon t(4) = -25.486, *p* = 1.42×10^-5^, diencephalon t(2.27) = - 10.954, *p* = 0.0052. N = 3 for each condition. Abbreviations: dpf - days post-fertilisation, KO - knockout, st - stage, WT - wild-type. Scale bars in b = 1 mm.

By 11 days post-fertilisation (dpf), *fgf8a* mutant embryos displayed severe disruptions across multiple body systems: axial shortening, abnormal tail curvature, dorsally displaced eyes and flattened neurocranium, particularly in the frontal and parietal regions (Fig. 3B), consistent with predicted pleiotropic effects of a major patterning gene. These external malformations coincided with a reduction in the dorso-lateral aspect of the optic tectum and anterior midbrain (white arrowhead in Fig. 3B dorsal view). Given that the cartilaginous cranial skeleton is not yet fully developed at this embryonic stage in cichlids (31), these defects are unlikely to arise from abnormal skeletal constraint, instead pointing to primary alterations in brain patterning or differential growth rates of the progenitor domains.

To dissect the neuroanatomical basis underlying these external phenotypic changes, we employed high-resolution μCT imaging. Volumetric analyses revealed significant reductions in the forebrain (diencephalon), midbrain (optic tectum) and hindbrain (rhombencephalon, including cerebellum) of *fgf8a* mutants compared to wild-type siblings (Fig. 3C). Notably, the diencephalon - one of the two forebrain regions enlarged in *A. calliptera* compared to *Rhamphochromis* - showed strong sensitivity to gene perturbation, directly implicating *fgf8a* in adult neuroanatomical differences between *A. calliptera* and *Rhamphochromis* (Fig. 1E).

### Variable transposable element insertions shape the regulatory landscape of the *fgf8a* locus in Malawi cichlids

Precise spatio-temporal control of vertebrate *fgf8* expression requires complex regulatory architectures comprising dozens of tissue-specific enhancers distributed across extensive genomic regions (23, 32). Given the pronounced expression differences between cichlid species (Fig. 2), and signatures of positive selection in the *fgf8a* regulatory region (Fig. 1, 15), we investigated whether *fgf8a cis*-regulatory evolution underlies brain morphological divergence between *A. calliptera* and *Rhamphochromis*.

Comparative analyses across African Great Lakes cichlids, other teleosts and non- teleost outgroups revealed conserved synteny surrounding *fgf8a*, universally flanked by *fbxw4* and *slc2a15a* (Fig. 4A, Supplementary Fig. 3). While *fgf8a* coding sequences exhibit high conservation across cichlids and other teleosts, the upstream regulatory landscape has been extensively restructured by transposable element (TE) insertions in Malawi cichlids.

**Figure 4.**
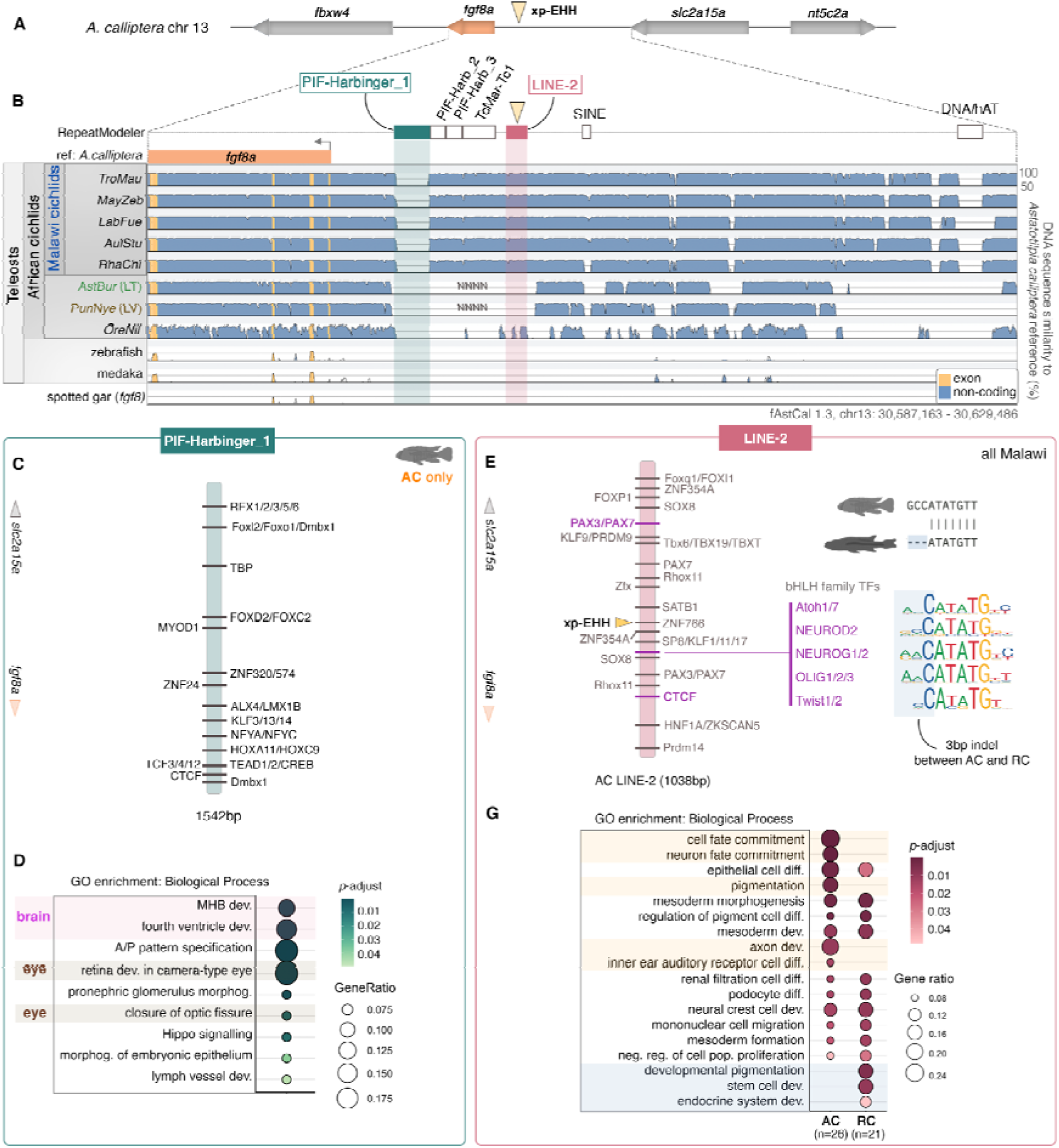
The genomic landscape upstream of *fgf8a* includes lineage-specific transposable element insertions with species-specific predicted regulatory potential in Malawi cichlids. **A.** Schematic of *fgf8a* locus in *A. calliptera*, including protein coding genes located ± 100 kbp from *fgf8a*. Arrows represent gene bodies and black lines represent non- coding intergenic regions. **B.** VISTA alignments comparing *fgf8a* and upstream non-coding region of *A. calliptera* with orthologous regions in model teleosts and *fgf8* of spotted gar (a non- teleost outgroup). Shading indicates regions of ≥100 bp that are ≥70% identical. Transposable element (TE) insertions from our curated library (see Methods) indicated by coloured boxes. Missing sequences in *Pundamilia nyereri* (PunNye 1.0) and *Astatotilapia burtoni* (AstBur 1.0) assemblies indicated by Ns. Extended version of this VISTA alignment with additional teleost species presented in Supplementary Fig. 3. **C.** Transcription factor binding sites (TFBSs) predicted for *A. calliptera*-specific PIF- Harbinger_1 (FIMO p-value < 0.0001, q-value < 0.05). **D.** Gene Ontology (GO) functional enrichment for Biological Processes based on the predicted TF binding motifs in PIF-Harbinger element (p < 0.05, FDR < 0.05). **E.** TFBSs predicted for the LINE-2 element of *A. calliptera* (FIMO p-value < 0.0001, q-value < 0.05). Sites with sequence variation between species highlighted in purple. Divergence in predicted TF motif repertoire between LINE-2 elements of *A. calliptera* and *Rhamphochromis* involves a shared site for multiple bHLH family TFs due to a 3 bp indel between the two species. **F.** Overlapping and divergent GO functional enrichment for Biological Processes of predicted TFs binding LINE-2 elements of *A. calliptera* and *Rhamphochromis*. GO terms highlighted in orange are unique to *A. calliptera*, terms highlighted in blue are specific to *Rhamphochromis*. Abbreviations: LT - Lake Tanganyika, LV - Lake Victoria, TFBM - transcription factor binding motif, TFBS - transcription factor binding site. AstBur - *Astatotilapia burtoni*, AulStu - *Aulonocara stuartgranti*, LabFue - *Labeotropheus fuelleborni*, MayZeb - *Maylandia zebra*, PunNye - *Pundamilia nyererei,* RhaChi - *Rhamphochromis* sp. “chilingali”, TroMau - *Tropheops* sp. “mauve”.

Within the ∼33kb intergenic region between *fgf8a* and *slc2a15a* in *A. calliptera,* we identified six distinct TE insertions (Fig. 4B): three full-length PIF-Harbinger DNA transposons, a Tc1/mariner element, a LINE-2 retrotransposon and a SINE, all located within 3-10kb upstream of the *fgf8a* coding sequence in *A. calliptera*, plus a putative non-autonomous DNA/hAT element (∼30.7 kb upstream of *fgf8a*) (Fig. 4B, Supplementary Table 2). Several insertions show lineage-restricted distributions: PIF-Harbinger_1 and hAT transposons were identified only in *A. calliptera*, whereas the SINE is absent from *Rhamphochromis* (Fig. 4B). These patterns suggest recent transposition events or lineage-specific deletions which might have created species-unique regulatory architectures at the *fgf8a* locus across the Malawi radiation. Fine-mapping of positive selection (xp-EHH, 15) revealed that the peak signal localises within the LINE-2 retrotransposon sequence (Fig. 4B), directly implicating this TE insertion in adaptive evolution and suggesting its functional integration into the *fgf8a cis-*regulatory landscape.

To assess whether TE-mediated regulatory evolution at *fgf8a* extends across the Lake Malawi radiation, we analysed TE insertion polymorphisms across the 120kb interval corresponding to the region of elevated xp-EHH values (Fig. 1C) in 15 cichlid species representing all major ecological groups (25). Phylogenetic reconstruction revealed that *Rhamphochromis* species harbour a significantly elevated frequency of private (clade-specific) TE insertions compared to all other Malawi lineages (Supplementary Fig. 3), indicating accelerated regulatory evolution at the *fgf8a* locus within the pelagic piscivore clade.

### Lineage-restricted transposable elements as candidate neurodevelopmental enhancers with species-specific regulatory functions

TEs are increasingly recognised as sources of novel enhancers co-opted into developmental gene networks (33). To test whether cichlid-specific TE insertions function as *fgf8a* regulatory elements, we analysed transcription factor (TF) binding motifs within each insertion. All identified TEs harboured multiple statistically significant TF binding sites (FIMO *p*-value < 10⁻ ^4^, q-value (FDR) < 0.05), consistent with regulatory competence (Supplementary Table 3).

We focused on two candidate elements with the strongest evidence for functional divergence: the LINE-2 element showing strong signatures of recent positive selection (xp-EHH signal) and the PIF-Harbinger_1 insertion present only in *A. calliptera* and located closest to *fgf8a* (Fig. 4B). Motif analysis (Supplementary Table 3) combined with Gene Ontology (GO) enrichment revealed that the PIF-Harbinger_1 element is enriched in binding motifs for TFs associated with brain development, including *lmx1b* (regulating *wnt1* and *fgf8* expression at midbrain-hindbrain boundary (34)), *dmbx1b* (35), TCF family members (Wnt pathway modulators (36) and RFX family factors involved in neural and sensory development (37–38) (Fig. 4C-D). These results suggest this insertion may integrate into established midbrain-hindbrain patterning circuits involving *fgf8a* and function as a novel developmental enhancer in neural morphogenesis specifically in *A. calliptera*.

The LINE-2 element, present across Malawi species (Fig. 4B), contained binding sites for key neurodevelopmental regulators: Pax7 (brain regionalisation, neural crest specification and muscle development (39–41)), Sp8 (involved in reciprocal regulation with *fgf8* (42)), and multiple bHLH (basic helix-loop-helix) family TFs involved in neurogenesis, lateral line development and craniofacial patterning (43–46) (Fig. 4E).

Detailed comparison of cloned LINE-2 sequences between *A. calliptera* and *Rhamphochromis* (Supplementary Fig. 5) revealed a species-specific difference with potential functional consequences: a 3-bp indel disrupting the binding site for multiple bHLH TFs in *Rhamphochromis* compared to a complete motif in *A. calliptera* (Fig. 4E-F). This *A. calliptera* LINE-2 is uniquely enriched for neuronal and mechanosensory pathways (e.g. axonogenesis, inner ear development), while the *Rhamphochromis* element is enriched for endocrine development (Fig. 4G).

Expression of several of the predicted TFs predicted to bind LINE-2 overlap spatially with the established *fgf8a* activity in teleosts, including *Pax7* at the midbrain-hindbrain boundary (39, 41), *Atoh7* in retinal ganglion cells (47) and *Twist2* in axial mesoderm (48). This spatial overlap supports a model where TE-derived enhancers, when bound by these TFs, could modulate *fgf8a* expression in specific developmental contexts. Additionally, the organisation of putative regulatory elements also differs between lineages due to species-specific insertions. In *Rhamphochromis*, the LINE-2 element is positioned closer to the *fgf8a* transcription start site compared to *A. calliptera* due to the intervening PIF-Harbinger_1 insertion, potentially further contributing to inter-specific differences in enhancer activity.

These findings suggest that TE insertions created lineage-specific regulatory modules near *fgf8a* that integrate multiple neural developmental pathways, with sequence evolution further fine-tuning their functions to contribute to species-specific neurodevelopmental programmes.

### *In vivo* enhancer assays of candidate TEs confirm regulatory functions with species-specific activity patterns in the nervous system

To test whether the cichlid-specific TE insertions function as developmental enhancers, we performed Tol2-mediated transgenic reporter assays of candidate cichlid TEs in a more tractable model, zebrafish (*Danio rerio*). We cloned the PIF- Harbinger_1 (specific to *A. calliptera*) and LINE-2 elements from both species upstream of an mCherry reporter gene and injected them into one-cell stage zebrafish embryos (Fig. 5A). Expression patterns were analysed in mosaic G0 embryos across multiple developmental stages (n > 80 embryos per construct across at least 2 independent experiments) (Supplementary Table 4).

**Figure 5.**
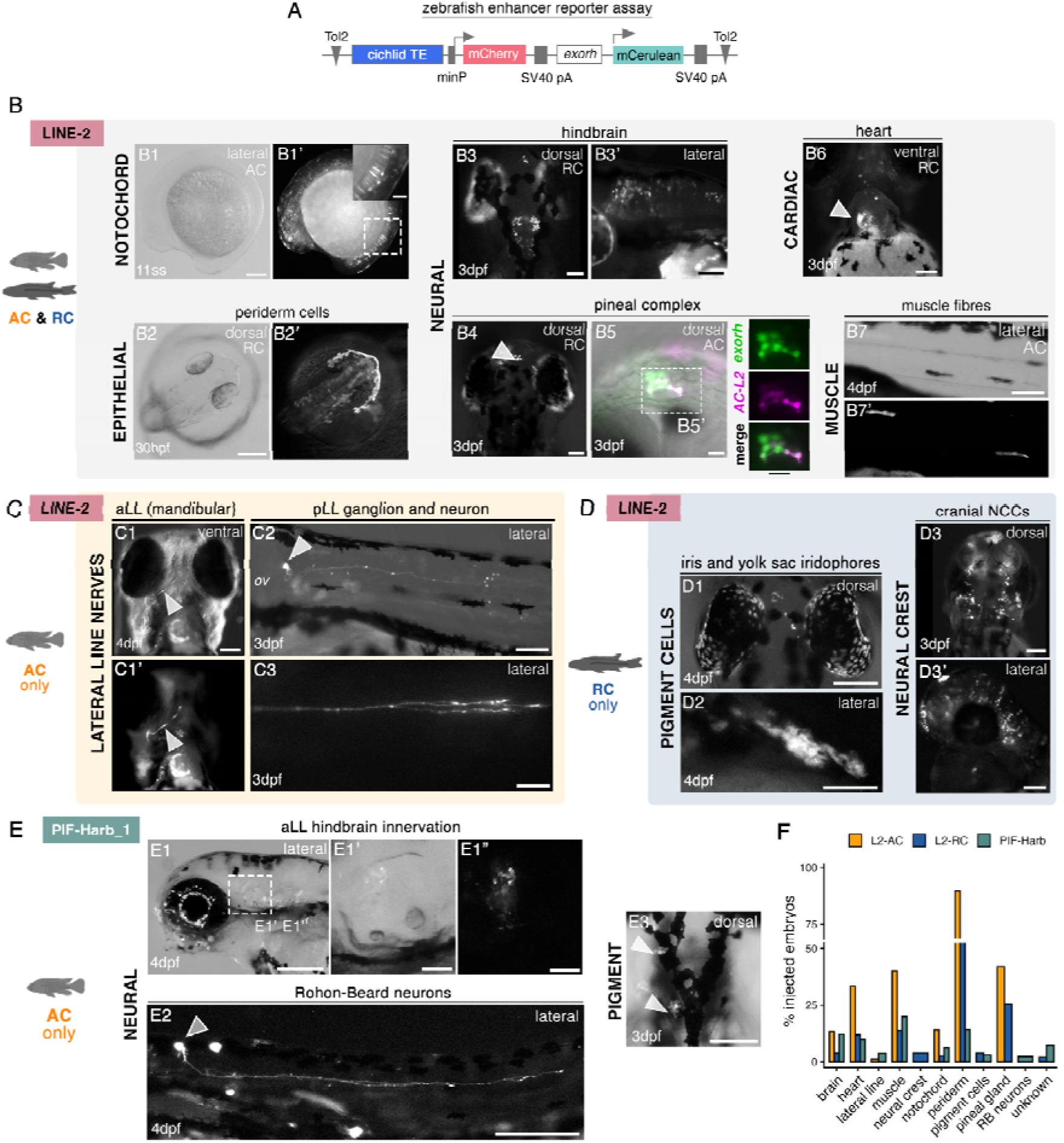
Transposable element insertions in *fgf8a* locus show enhancer activity *in vivo* with species-specific outcomes associated with neural, sensory and neural crest developmental programmes. **A.** Schematic of the zebrafish enhancer reporter assay. Cichlid transposable element sequences (LINE-2 or PIF-Harbinger_1) were cloned upstream of a mouse minimal promoter (minP) driving mCherry reporter expression in a modified Tol2-based construct (Kemmler et al. 2024). A constitutive *exorh* promoter drives mCerulean as a transgenesis control. The constructs were injected into one-cell stage zebrafish embryos for assessment of tissue-specific enhancer activity in mosaic animals. **B.** Shared LINE-2 enhancer activities in both species. Expression in notochord (B1- B1’), periderm (B2-B2’), hindbrain (B3-B3’), pineal complex (B4-B5’), heart (B6) and muscle fibers (B7-B7’) at indicated developmental stages. Arrowheads indicate specific expression domains. **C.** *A. calliptera* LINE-2-specific lateral line expression. Anterior lateral line mandibular nerve (C1, arrowhead) and posterior lateral line ganglion and neurons (C2-C3, arrowheads) at 3-4 dpf. Expression absent in *Rhamphochromis* LINE-2 constructs. Arrowhead in (C2) indicates the pLL ganglion. **D.** *Rhamphochromis* LINE-2-specific expression in neural crest derivatives. Cranial neural crest cells (D3-D3’) and pigment cells including iris and yolk sac iridophores (D1-D2) at 3-4 dpf. Expression not detected in *A. calliptera* LINE-2 constructs. **E.** PIF-Harbinger_1 expression (*A. calliptera*-specific element). Expression in the cells of the rhombencephalic nuclei (E1’), Rohon-Beard sensory neurons in the spinal cord (E2, arrowhead) and pigment cells (E3) at 3-4 dpf. Dashed box in (E1) indicates region shown at higher magnification in (E1’) and (E1’’). **F.** Quantification of enhancer activity across tissue types. Data represent percentage of mCherry-positive embryos showing expression in each tissue (n > 80 embryos per construct across three independent experiments). All images show representative examples of mosaic G0 embryos. Abbreviations: AC, *Astatotilapia calliptera*; aLL, anterior lateral line; dpf, days post-fertilisation; hpf, hours post-fertilisation; ov, otic vesicle; pLL, posterior lateral line; RC, *Rhamphochromis* sp. ‘chilingali’. Scale bars equal to 250μm in E1-E2, 200μm in B1, B2, B5, C3, 100μm in B3, B4, B6, B7, C1, C2, D1-D3’, E1’, E3, 50μm in B1’ and 20μm in B5’.

All three TE constructs drove robust, tissue-specific reporter expression, demonstrating their capacity to function as *bona fide* developmental enhancers (Fig. 5B-F). This included shared expression in the periderm and muscle tissues, with quantified proportions shown in Fig. 5F. Importantly, enhancer activities showed marked bias towards neural and sensory tissues, aligning with both the predicted transcription factor binding profiles and established roles of fgf8a signalling in the development of these systems.

Both LINE-2 variants showed shared activities in the hindbrain and pineal gland (Fig. 5B, B3-B5’), the latter directly relevant to the diencephalic differences observed between adult *A. calliptera* and *Rhamphochromis*. In addition, species-specific differences emerged: only the *A. calliptera* LINE-2 drove reporter expression in the lateral line system (anterior and posterior lateral line neurons and ganglion, Fig. 5C, C1-C3). This appears consistent with the binding sites prediction in *A. calliptera* for Atoh1 and Neurod in these species, as these are key regulators of mechanosensory hair cell development and innervation in the lateral line (43–44). In contrast, the *Rhamphochromis* LINE-2 was active in cranial neural crest cells and pigment derivatives, including iris and yolk sac iridophores (Fig. 5D, D1-D3’).

The *A. calliptera*-specific PIF-Harbinger_1 showed extensive activity within the nervous system (Fig. 5E), including inner ear lateral line innervation into the hindbrain (Fig. 5E, E1-E1”), Rohon-Beard sensory neurons in the spinal cord (Fig. 5E, E2) and pigment cells (Fig. 5E, E3).

Critically, several enhancer activities overlapped with known *fgf8a* expression domains in zebrafish, particularly the pineal gland, suggesting these TEs could directly modulate *fgf8a* expression in their native cichlid contexts in a species-specific manner. However, additional activities in lateral line and neural crest suggest these elements may also regulate other target genes or have been co-opted for novel functions. These findings demonstrate that TE evolution has generated species-specific enhancer toolkits that are likely to function in the neural and sensory systems that show morphological divergence between cichlid species, providing a mechanistic link between regulatory innovation and adaptive nervous system evolution.

## Discussion

The rapid diversification of brain and sensory systems during adaptive radiation has long captivated evolutionary neurobiologists, yet the underlying molecular mechanisms remain incompletely understood. Here we present converging lines of evidence linking *fgf8a* regulatory evolution to neuroanatomical diversification in ecologically divergent Lake Malawi cichlids *Astatotilapia calliptera* and *Rhamphochromis* sp. ‘chilingali’. First, these species exhibit striking differences in forebrain domain proportions, implicating mosaic brain evolution between ecologically divergent species. Second, these morphological differences correlate with species-specific *fgf8a* expression patterns during critical developmental windows. Third, regulatory analysis of *fgf8a* locus reveals species-specific TE-derived enhancer elements with divergent activities in neural and sensory systems. Together, these findings demonstrate how modifications to a conserved developmental programme can generate neuroanatomical diversity.

### Expression and functional evidence implicate *fgf8a* in cichlid brain diversification

Complementary expression and functional analyses provide compelling evidence for *fgf8a*’s role in cichlid neural evolution. Species-specific *fgf8a* expression differences emerge early and persist through brain development in territories that correspond to adult morphological variation between *A. calliptera* and *Rhamphochromis*. The apparent mismatch between embryonic expression patterns and adult brain sizes, where *Rhamphochromis* exhibits stronger forebrain *fgf8a* expression, yet smaller forebrain volumes in adulthood, highlights the complexity of morphogen-mediated patterning. Early presumptive forebrain expression in *A. calliptera* may establish developmental trajectories that persist despite later downregulation, consistent with studies showing *fgf8a* protein gradients influence territories beyond immediate expression domains during zebrafish gastrulation (50). Alternatively, species-specific receptor expression patterns may modulate morphogen sensitivity, emphasising the need for future *fgf8a* protein gradient and receptor mapping studies across cichlid development.

CRISPR/Cas9-mediated knockdown in *A. calliptera* confirmed *fgf8a*’s essential role for the development of multiple forebrain, midbrain, and hindbrain territories in cichlids, consistent with its pleiotropic role in vertebrate neurodevelopment (11,17). While we did not expect knockout phenotypes to directly replicate the natural morphological differences between *A. calliptera* and *Rhamphochromis*, we also observed phenotypic outcomes diverging from the zebrafish *ace* mutants, including optic tectum reduction without telencephalic volume loss (11, 51, 52), suggesting cichlid-specific developmental outcomes. Our morphological analysis revealed modular evolution of brain regions between *A. calliptera* and *Rhamphochromis*, where diencephalon and telencephalon volumes evolved independently of other structures, consistent with *fgf8a*’s region-specific expression and functional effects.

Taken together, our results demonstrate that modifications to *fgf8a* expression patterns, signal strength and distribution contribute to the evolution of neural diversity between these ecologically distinct Lake Malawi species. The broad phenotypic effects observed in CRISPR mosaic mutants, affecting regions beyond our focal *A. calliptera* versus *Rhamphochromis* comparison, suggest *fgf8a* regulatory modifications could potentially underlie brain morphological differences across the broader cichlid radiation.

### Transposable element insertions provide regulatory mechanisms for nervous system diversification

Despite strong syntenic conservation across teleosts, the cichlid *fgf8a* locus has accumulated numerous TE insertions. These fall into two categories: (i) unique insertions like PIF-Harbinger_1 or hAT in *A. calliptera* potentially introducing lineage-restricted novel regulatory modules, and (ii) shared elements like LINE-2 that have undergone sequence divergence, resulting in distinct regulatory potentials between species. The latter parallels snake limb evolution, where a single indel in *shh* enhancer directly causes limb loss (53), demonstrating how small regulatory changes can drive major morphological adaptations.

Zebrafish enhancer assays confirmed that TE-derived sequences function as developmental enhancers with pronounced neural and sensory system activity. Critically, both *A. calliptera* and *Rhamphochromis* LINE-2 elements showed robust expression in the pineal gland, a structure central to ecologically relevant circadian behaviours and a part of the diencephalon that differed in the adult volume between species. Shared enhancer activity in this structure therefore points to LINE-2-specific regulatory role in development of the photoreceptive circuitry in cichlids, with potential contribution of this structure to gross volumetric differences involving this forebrain region.

Our results further point to potential regulatory integration across divergent neural systems, as TE-derived enhancers drove expression in brain territories and lateral line sensory system - both traits that differ between *A. calliptera* and *Rhamphochromis* (54). However, zebrafish assays cannot fully recapitulate native cichlid developmental contexts, and the observed enhancer activities may thus not fully capture species-specific differences that would be evident in the native cichlid system. Moreover, the potential combinatorial activity of multiple regulatory elements require validation in cichlid embryos to determine whether these elements collectively drive the precise expression differences underlying natural brain variation between these species.

### Modular regulatory architecture at *fgf8a* enables both mosaic and integrated evolutionary innovation

The Fgf developmental programme is both highly conserved and dose-dependent (55), making it ideal for evolutionary fine-tuning via modular regulatory changes. TE insertions provide a rapid mechanism for introducing regulatory elements without disrupting essential functions (56), enabling developmental stability to coexist with morphological innovation. Our findings suggest that this modular architecture at cichlid *fgf8a* may facilitate evolution through two complementary mechanisms.

First, individual TE insertions can create discrete enhancer units that drive expression in specific developmental contexts, as demonstrated by our functional validation showing spatially restricted activities. Such tissue-specific enhancers could evolve independently, influencing their targets to evolve in distinct directions, akin to the mosaic brain evolution we documented between *A. calliptera* and *Rhamphochromis*. The species-specific differences in TE insertion repertoires we identified - with *A. calliptera* possessing unique PIF-Harbinger and hAT elements - further support this modular evolutionary mechanism by introducing lineage-restricted regulatory potential.

Conversely, the tight genomic clustering of multiple regulatory elements with the *fgf8a* locus creates potential for coordinated evolution through linkage. This could be further enhanced given *fgf8a*’s pleiotropic developmental functions, with regulatory changes affecting this single gene potentially simultaneously influencing multiple trait systems. Our observation that TE-derived enhancers drive expression in brain territories, sensory systems, and neural crest derivatives - all of which show phenotypic variation between our focal species - suggests this integrated regulatory evolution could facilitate correlated adaptive responses across functionally related systems.

The balance between these evolutionary mechanisms may be influenced by enhancer redundancy, as demonstrated in mouse *fgf8* where overlapping regulatory elements provide developmental robustness while enabling evolutionary flexibility (57). Our phylogenetic analysis supports ongoing regulatory evolution at multiple scales, with lineage-specific insertions in *A. calliptera*, coupled with elevated TE polymorphism in *Rhamphochromis* spp., indicate accelerated evolution in this piscivorous clade. Notably, the strong xp-EHH signal in this locus likely reflects reduced variation in A. calliptera and other clades relative to the highly polymorphic *Rhamphochromis*, rather than necessarily indicating *A. calliptera*-specific positive selection. The persistence of multiple TE insertions and sequence variants across the radiation, combined with our functional evidence, suggests this locus maintains substantial regulatory potential for generating phenotypic variation affecting multiple trait systems - including brain architectures and sensory development - underlying ecological specialisation across different cichlid lineages.

The modular regulatory architecture - combining conserved developmental functions with lineage-specific elements and sequence variation - provides a mechanistic framework for understanding how adaptive radiations might balance essential developmental requirements with morphological innovation. Future experimental validation in native cichlid developmental contexts will be essential to establishing the specific adaptive contributions of individual *fgf8a* regulatory elements.

### Implications for understanding mechanisms of brain evolution during adaptive radiations

Our findings position *fgf8a* within a broader network of developmental pathways previously implicated in cichlid brain evolution, including Wnt and Hedgehog (8–10). Given the extensive cross-regulation among Fgf, Wnt, Bmp and Hedgehog signalling that controls forebrain and midbrain-hindbrain boundary patterning (58–60), *fgf8a* represents an additional axis for evolutionary modification. This multi-dimensional patterning system facilitates rapid evolutionary fine-tuning, where subtle spatiotemporal shifts in any component can yield substantial neuroanatomical changes - as exemplified by the mosaic evolution of brain regions in ecologically divergent species we documented in this study. The spatially restricted expression patterns we observed for TE-derived enhancers suggest that regulatory modifications can achieve region-specific effects within this network, enabling the independent evolution of different brain territories while maintaining overall developmental coordination. Understanding whether regulatory changes across Fgf, Wnt, and other signalling pathways evolve co-ordinately or independently represents an important avenue for future research into the mechanisms driving adaptive brain diversification.

More broadly, while TE-mediated regulatory evolution has been implicated in diverse adaptive traits - from mammalian placenta development (61) to stickleback armour reduction (62) and cichlid egg-spots (63) - direct functional links to neural diversification have remained rare. Our study demonstrates how TE-derived regulatory modifications can integrate into conserved developmental programmes to generate heritable differences in brain and sensory architecture. Such regulatory flexibility may be particularly advantageous for adaptive radiations, where coordinated evolution of multiple trait systems enables rapid ecological specialisation.

The integration of our experimental validation with phylogenetic evidence for ongoing regulatory evolution provides insights into how genomic and developmental constraints can be overcome during rapid morphological diversification. However, establishing the specific contributions of these regulatory innovations to natural phenotypic variation represents the critical next step for understanding the role of regulatory evolution in adaptive brain diversification.

## Conclusions

Our study demonstrates that *fgf8a* expression and functional evolution contribute directly to cichlid brain diversification, while regulatory innovations through transposable element insertions provide additional evolutionary potential. The cichlid system demonstrates that conserved developmental programs can accommodate extensive regulatory modification to generate adaptive neural diversity. Through both expression evolution and regulatory innovation via transposable elements, *fgf8a* exemplifies how master developmental genes serve as evolutionary targets for brain morphological diversification to facilitate mosaic evolution. Understanding these mechanisms provides fundamental insights into the genetic and neurodevelopmental bases underlying nature’s most spectacular adaptive radiations.

## Methods

### Animal husbandry and embryo collection

Adult cichlid stocks were maintained at the Department of Zoology, University of Cambridge (UK) fish facility in accordance with UK Home Office regulations. Cichlid embryos were collected, cultured, staged and imaged as described previously (64).

Eggs were removed from mouthbrooding females and maintained in tap water in 6- well plates (Corning) on a slowly moving orbital shaker at 27°C.

The maintenance of adult zebrafish was conducted in accordance with the Animals (Scientific Procedures) Act 1986 Amendment Regulations 2012, following ethical review by the University of Cambridge Animal Welfare and Ethical Review Body (AWERB) at the University Biomedical Service Aquatics facility at the Department of Physiology and Neuroscience, University of Cambridge (UK). AB and Tupfel long fin (TL) wild-type strains were used. All embryos were raised in standard E3 media supplemented with 0.0001% methylene blue at 27°C and staged according to ref. 65.

### Adult images

Adult fish were photographed using a Sony α6600 camera with Sony E 30mm f/3.5 lens.

### Brain micro- and nano- computer tomography (CT) scans

We used diffusible iodine-based contrast-enhanced computed tomography (Dice-CT) to measure and compare the volumes of multiple higher-level brain structures between adult *Rhamphochromis* sp. ‘chilingali’ and *Astatotilapia calliptera*, as well as between wild-type and CRISPR-knockout embryos of *A. calliptera*. For adults, whole fish were stained for a minimum of 14 days in a 1% iodine, ethanol solution. This solution was refreshed around every 7 days. Prior to CT scanning, fish were put into 70% ethanol ready to be scanned. For embryos, samples were stained overnight in a 1% iodine ethanol solution, before being washed in 70% ethanol, and immobilised in 5% agar to prevent movement during scanning.

All samples were imaged using a Nikon XTH 225ST X-ray tomography scanner (University of Bristol, UK). For adults, there was a maximum of 8 adult fish per scan, arranged radially. Scans were taken with a resolution of approximately 30µm, an exposure of 708ms, 3141 projections and 1 frame per projection. Embryos were scanned one at a time, with a resolution of approximately 3µm, an exposure of 1415ms, 3141 projections and 1 frame per projection. Volumes were reconstructed using Nikon μCT software and VG Studio Max and exported as an image stack of Tiff files. We then cropped these multi-fish image stacks using the Fiji/ImageJ (66) plugin Crop (3D), to generate a separate stack for each sample. Dragonfly (Comet) was used to implement Contrast Limited Adaptive Histogram Equalization (CLAHE), and to then carry out segmentation and structure volume measurement. The volume of a total of five major structures was measured: the brain stem (allometric control), diencephalon, rhombencephalon, optic tectum and telencephalon.

### Whole mount *in situ* hybridization by Hybridization Chain Reaction (HCR)

Cichlid embryo dissection, fixation and HCR were conducted as described previously (Marconi, Vernaz et al., 2024). In brief, embryos were dissected and fixed in 4% paraformaldehyde (PFA) in phosphate buffered saline (PBS) (pH 7.4) at 4°C overnight. Specimens were stepwise dehydrated into 100% methanol and stored at - 20°C under further analyses. Embryos were rehydrated and stained with v3 HCR (30) modified for cichlid embryos (15) with probes designed against *A. calliptera* genome sequences (fAstCal1.3, Ensembl 111). Probe sets (14-20 pairs per gene) were purchased from Molecular Instruments (Supplementary Table 5) and verified by BLAST analysis.

Following HCR staining, specimens were counterstained with 10 nM DAPI in 5xSSCT at 4°C overnight. Samples were cleared with DEEP-Clear solution 1.1 (67) for 15 min at room temperature, washed twice with 5x SSCT and mounted on microscopy slides (ThermoFisher Scientific) with #1.5 coverslips (Corning) in 25% Vectashield (Vector) - 75% (glycerol:Tris-HCl pH 7.5) mounting solution. Clear nail varnish was used to seal the edges of the slide and all samples were cured overnight at 4°C protected from light before imaging on inverted confocal microscope Olympus FV3000 at the Imaging facility of the Department of Zoology, University of Cambridge (UK). All *in situ* hybridization experiments were performed with multiple specimens from different clutches (at least 3 individuals per clutch, repeated at least once with specimens from alternative clutches) to fully characterise the expression patterns. HCR images were processed using ImageJ/Fiji (66) and 3D reconstructions generated using Imaris software (Bitplane). Volumetric measurements were obtained from gene expression domains using HCR signal and for total brains using DAPI signal to outline the regions of interest. Measured volumes of gene expression and total brain were averaged over at least 3 replicates.

### CRISPR/Cas9 genome editing

CRISPR/Cas9-mediated transgenesis in *Astatotilapia calliptera* was performed as described previously (15). Briefly, two sgRNAs targeting exon 2 of *fgf8a* (ENSACLG00000037698) (sgRNA-513 5’-ACCGTGCGCATCTCCATCCT-3’ and sgRNA-522 5’-GGTAGATCCGGATCAACCTG-3’) were designed using Geneious Prime (Dotmatics) software and verified for off-target activity against the *A. calliptera* 1.3 genome (Ensembl 113). sgRNAs were purchased from Integrated DNA Technologies (ITD) as Alt-R CRISPR-Cas9 sgRNA (2 nmol).

One-cell stage embryos were microinjected with a mix of two sgRNAs (at 150 ng/μl each), Texas Red (0.5% v/v) and Alt-R™ S.p. HiFi Cas9 Nuclease V3 (IDT, 100 ng/μl).

Embryos were imaged at 11 dpf using a Leica M165FC stereoscope with a DFC7000T camera under reflected light brightfield. All specimens were positioned in 1% low melting point agarose (Promega) and anesthetized with 0.02% MS-222 (Sigma-Aldrich) to immobilise during imaging.

Genomic DNA was extracted from specimens sacrificed by overdose of 0.5% MS-222 (Sigma-Aldrich) using PCRBIO Rapid Extract Lysis Kit (PCRBiosystems). Fragments of 470 bp surrounding sgRNA target sites were amplified using PCRBIO HS Taq Mix Red (PCRBiosystems) with an annealing temperature of 55°C using following primer pairs: 5’-CTGAAATCTACCTAAATGAACGCAG-3’ and 5’- ACCACCCCAAAACACAAAC-3’. Amplicons were purified with QIAquick PCR Purification Kit (Qiagen) before Sanger sequencing using the same primers. All protocols were conducted following manufacturer’s instructions. CRISPR edit sites were inferred using the Synthego ICE CRISPR analysis tool (https://ice.synthego.com/).

### Genomic analyses

**Transposable element annotation.** We followed the protocol of ref. 68 with modifications. An initial *de novo* library was generated using RepeatModeler 2.0.3, with the option -genomeSampleSizeMax 900000000 to sample the whole genome, yielding 2,955 candidate TEs. Unlike the original protocol, instead of producing a priority list for manual curation, we applied a *de novo* extension method to recover full TE sequences, implemented later in MCHelper (69).

Sequences were clustered into families using CD-HIT 4.8.1 using parameters -aS 0.8 -c 0.8 -G 0 -g 1 -b 500, following the ‘80-80-80 framework’ of ref. 70. Structural features (LTRs, TIRs and polyA sequences) were identified, and their ORFs and protein products were verified. Classification was informed by similarity searches against RepBase (via BLAST 2.12.0) and RepeatClassifier 2.0.3, with incomplete elements removed from families with well-defined structures. Large palindromes or elements with TIRs were retained as putative non-autonomous DNA transposons, but not assigned to families.

To further enrich the library, we combined those results with those from a preliminary version of Pantera (71), run on a pangenome built from a pggb alignment of the genomes of *Astatotilapia calliptera* and *Maylandia zebra*. The final library comprised 463 TE families (MWCI3.2) and was deposited in Dfam (72).

The fAstCal1.2 genome was annotated with the most recent version of 626 families (with LTRs divided into internal and LTR components) using RepeatMasker version 4.1.2-p1. Mobile element (ME) genotyping was performed with MEGAnE (73), which infers ME polymorphisms from split and discordant short reads relative to a reference. We ran MEGAnE with options -no_sex_chr -lowdep -skip_unmapped on 2,207 samples of cichlids from Lake Malawi (25), all mapped to fAstCal1.2 (RefSeq:GCF_900246225.1).

### Comparative genomic analysis using VISTA

Comparative sequence analysis of the *fgf8a* locus was performed using the VISTA (https://genome.lbl.gov/vista/index.shtml) to identify conserved regions across teleost species. The analysis focused on a ∼40 kbp genomic interval centred on *fgf8a* and its upstream regulatory region in *Astatotilapia calliptera* (assembly fAstCal1.3, Ensembl 111).

For comparative alignment, we extracted orthologous genomic sequences from cichlid species including *Aulonocara stuartgranti*, *Tropheops* sp. ‘mauve’ and *Rhamphochromis* sp. ‘chilingali’, *Maylandia zebra*, *Pundamilia nyererei*, *Astatotilapia burtoni*, and model organisms *Danio rerio* (zebrafish) and *Oryzias latipes* (medaka), as well as the non-teleost outgroup *Lepisosteus oculatus* (spotted gar) from publicly available genome assemblies on Ensembl (111) and NCBI. Sequences for *Labeotropheus fuelleborni* were extracted from an unpublished assembly generated by Richard Durbin Lab (University of Cambridge, UK) available upon request. In all cases, genomic coordinates for *fgf8a* orthologs were identified through synteny analysis and confirmed by BLAST searches against respective genome assemblies.

VISTA alignments were generated using the following parameters: minimum sequence length of 50 bp and minimum sequence identity threshold of 70% for identifying conserved regions with the *A. calliptera* sequence as the reference genome for all comparisons. The VISTA alignment of the same region with additional teleost species is provided in Supplementary Figure 4.

**Transcription factor binding analysis.** TE sequences located between *fgf8a* and *slc2a15a* were scanned for transcription factor (TF) binding sites using FIMO (Grant et al., 2011) and JASPAR 2024 vertebrate motifs (75) (p < 0.0001, q-value (FDR) < 0.05). Gene Ontology enrichment analysis was performed using clusterProfiler (76) with significance thresholds of p <0.01 and FDR <0.05. Zebrafish orthologues of cichlid genes were used (Supplementary Table 6) and, where appropriate, all existing zebrafish paralogues were included.

**TE polymorphism and phylogenetic analysis.** To investigate TE polymorphisms in *fgf8a* locus across Lake Malawi cichlid populations, we analyzed 180 samples representing 15 distinct populations of species using MEGAnE (73) in 180 samples from 15 different species populations of Lake Malawi. We quantified TE insertions that were private to *Rhamphochromis* spp. (i.e., present only in this clade and absent in other taxa) compared to insertions shared with or exclusive to other species. Private TE insertions were operationally defined as those present in a focal species but absent in all sister species, indicating insertion events likely occurring after speciation. The analysis was performed using a 119,968 base pair sliding window on chromosome 13. This window encompassed the complete coding sequences of *fgf8a* and *slc2a15a*, flanked by 20 kb upstream of *slc2a15a* and 20 kb downstream of *fgf8a*, capturing the regulatory landscape surrounding these genes with elevated xp-EHH values. We performed sliding window analysis across this genomic region to identify the distribution of private TE insertions in *Rhamphochromis* relative to other Lake Malawi cichlid species. A Neighbour-Joining phylogenetic tree based on TE insertion polymorphism patterns was constructed using the ape package (v5.0) in R (77). This approach allowed us to assess the phylogenetic signal of TE-based variation and the distinctiveness of the *Rhamphochromis* lineage within the Lake Malawi radiation.

### Tol2 enhancer reporter assays

Transposable element sequences were cloned upstream of a minimal promoter driving mCherry reporter expression in a modified Tol2 construct (Kemmler et al. 2024). Genomic DNA was extracted from fin clips of adult wild-type laboratory stocks of *Astatotilapia calliptera* and *Rhamphochromis* sp. “chilingali” using the Zymo DNA- miniprep kit according to the manufacturer’s instructions. LINE-2 elements were PCR-amplified from genomic DNA using Phusion Green Master Mix (ThermoFisher) following a standard PCR protocol and the primers containing 5’ overhangs (in italics) corresponding to restriction enzyme cut sites: forward 5’- *GTCAGC*TAGCACCACACAGGGACCTTGAAA-3’ and reverse 5’- *GTCAGTCGAC*GAGTCAGGTACTTCTGATTAGAGACCTT-3’. Due to the highly repetitive nature of these elements, which rendered specific amplification challenging, we included an additional 200 bp upstream of each LINE-2 element that did not overlap with other TEs and maintained homologous regions between *A. calliptera* and *Rhamphochromis*. PCR products were analysed by gel electrophoresis to confirm expected sizes and their sequence validated by Sanger sequencing.

The sequence of PIF-Harbinger_1 was extracted from *A. calliptera* genome assembly (AstCal1.3) and synthesised by GENEWIZ with restriction enzyme overhangs and supplied in pUC-GW-Amp plasmid. The insert was excised using NheI and SalI FastDigest restriction enzymes (ThermoFisher) in 20 μl reactions for 3h at 37°C, followed by thermal inactivation of restriction enzymes at 65°C for 15 mins. The backbone Tol2 plasmid (pCK082 desmaMCS:minprommCherry,exorh:mCerulean) was obtained from the Mosimann Lab (University of Colorado, USA) via AddGene (product code #200015) and verified by restriction digestion with NheI and SalI FastDigest enzymes. The *desma* sequence was excised by restriction enzyme digestion and gel electrophoresis, and the remaining plasmid backbone was gel-purified using the Qiagen gel extraction kit.

Elements were ligated into a linearised backbone vector using 0.2 μl of T4 ligase (ThermoFisher) at optimised insert:vector ratios (3:1 for LINE-2, 5:1 for PIF-Harbinger_1) in 20 μl reactions. Ligation reactions were incubated for 30 minutes at room temperature followed by overnight incubation at 4°C. Ligated products were transformed into *E. coli* DH10B Max Efficiency cells (ThermoFisher) using standard chemical transformation protocol. Transformed cells were plated on LB agar plates containing ampicillin (100 μg/ml) and incubated overnight at 37°C.

Colonies were initially verified by colony PCR on lysed bacterial cells using the original cloning primers as well as M13 reverse primer (5’- CAGGAAACAGCTATGAC-3’) and a custom mCherry primer (5’- TTGGTCACCTTCAGCTTGG-3’) to confirm both the presence and orientation of inserts. All plasmids were sequenced using next-generation sequencing by Plasmidsaurus (GenBank files are provided as Supplementary Files).

To synthesise Tol2 transposase mRNA *in vitro*, the NLS-Transposase-NLS-pA- pT3TS-Dest plasmid was linearised with the restriction enzyme XhoI and purified by phenol:chloroform extraction, followed by ethanol precipitation. The resulting DNA was resuspended in nuclease-free Milli-Q water. Capped mRNA was transcribed in vitro using the mMESSAGE mMACHINE T3 Transcription Kit (Thermo Fisher Scientific, Cat. No. AM1348) with 1 µg of linearised plasmid as template. One-cell stage zebrafish embryos were injected with 1-2 nl of injection mix containing 25 ng/μl plasmid DNA and transposase mRNA. Embryos were maintained at 27°C in E3 medium, anaesthetised with 0.016% Tricaine-S (MS-222, ThermoFisher) and imaged using either Leica M165FC stereoscope with a DFC7000T camera or Zeiss Axio Observer widefield microscope using Zeiss Plan-Apochromat 40x (1.4NA) oil immersion objective and appropriate fluorescence filters.

### Statistical analysis

Statistical analyses were performed using R (v3.6.03) in R Studio software (v2025.09.0+387). Specific tests used are indicated in main text and figure legends. For morphological comparisons, one-way ANOVA was used with Tukey’s HSD for post-hoc pairwise comparisons. Student’s t-tests were applied for expression domain comparisons.

### Data and code availability

All scripts used to analyse the data and generate figures presented in this manuscript can be accessed at: https://github.com/Santos-cichlids/fgf8a-signalling-Malawi-cichlid-brain-divergence.

## Acknowledgements

We thank Moritz Blumer for providing genomic resources and Carlos Camacho de la Macorra for sharing Tol2 mRNA. We are also grateful to the technicians at the Department of Zoology for facility maintenance and fish husbandry. This work was supported by funding from the Natural Environment Research Council (NERC).

## Authors’ contributions

AM, SHM and MES devised the study and analyses. AM collected adult material, conducted CRISPR/Cas9 mutagenesis, embryo imaging, phenotyping, DNA collection, Tol2 construct assembly, HCR staining, imaging and data processing. AM and DS performed transgenic enhancer assays. JM conducted specimen staining, CT scanning and brain segmentation. PS collated TE library and performed data analysis. JE performed data analysis. RD contributed to whole-genome data collection. BS contributed zebrafish stocks for enhancer assay analysis. AM, SHM and MES wrote and revised the manuscript. All authors read and approved the final manuscript.

