## Supplementary Figures for "fgf8a signalling shapes brain divergence between Malawi cichlids"

Supplementary Information for *fgf8a signalling shapes brain divergence between Malawi cichlids* (Marconi et al.)

**Supplementary Figures**

**Supplementary Figure 1:** Additional brain morphometry data for brain structures not shown in Figure 1. Rhombencephalon and optic tecta volumes do not show significant allometric shifts between *A. calliptera* (AC) and *Rhamphochromis* sp. ‘chillingali’ (RC).

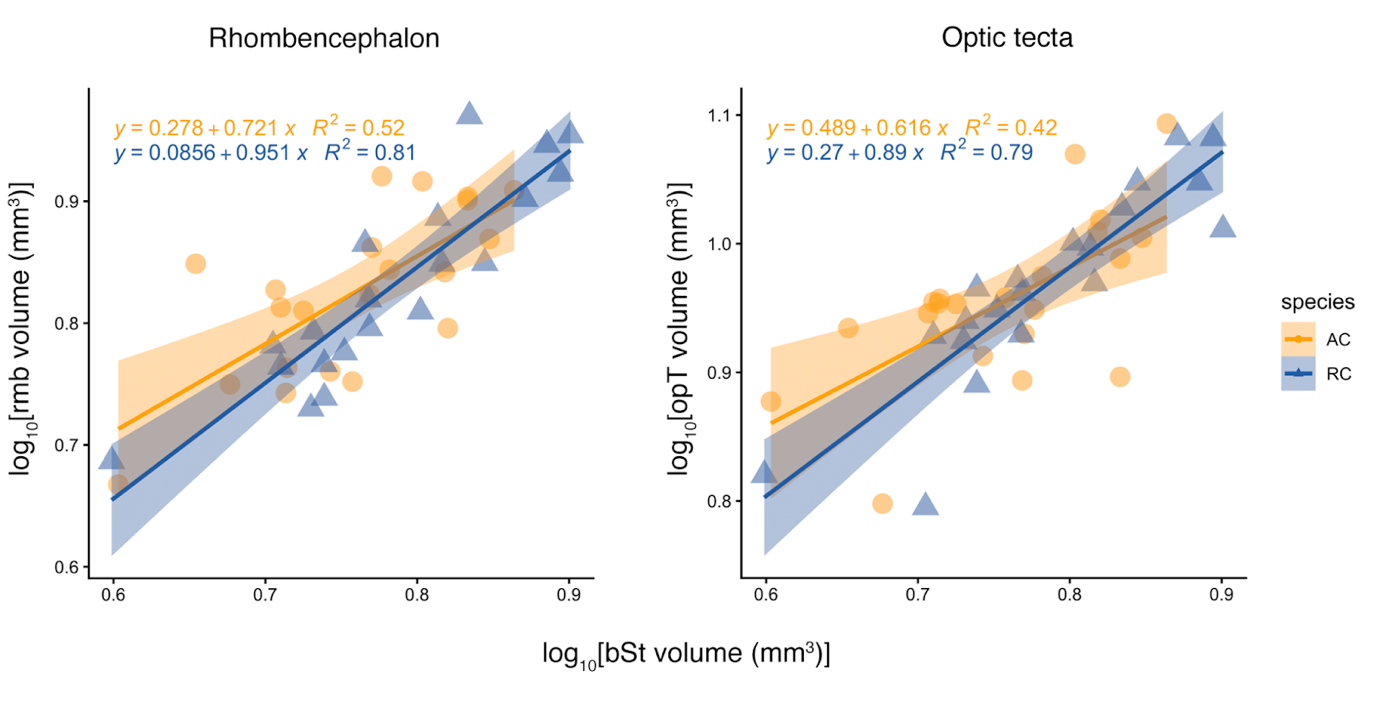

**Supplementary Figure 2:** CRISPR/Cas9 knockout Sanger sequencing validation. Example Synthego ICE analysis graphs showing CRISPR editing efficiency in exon 2 of *fgf8a*.

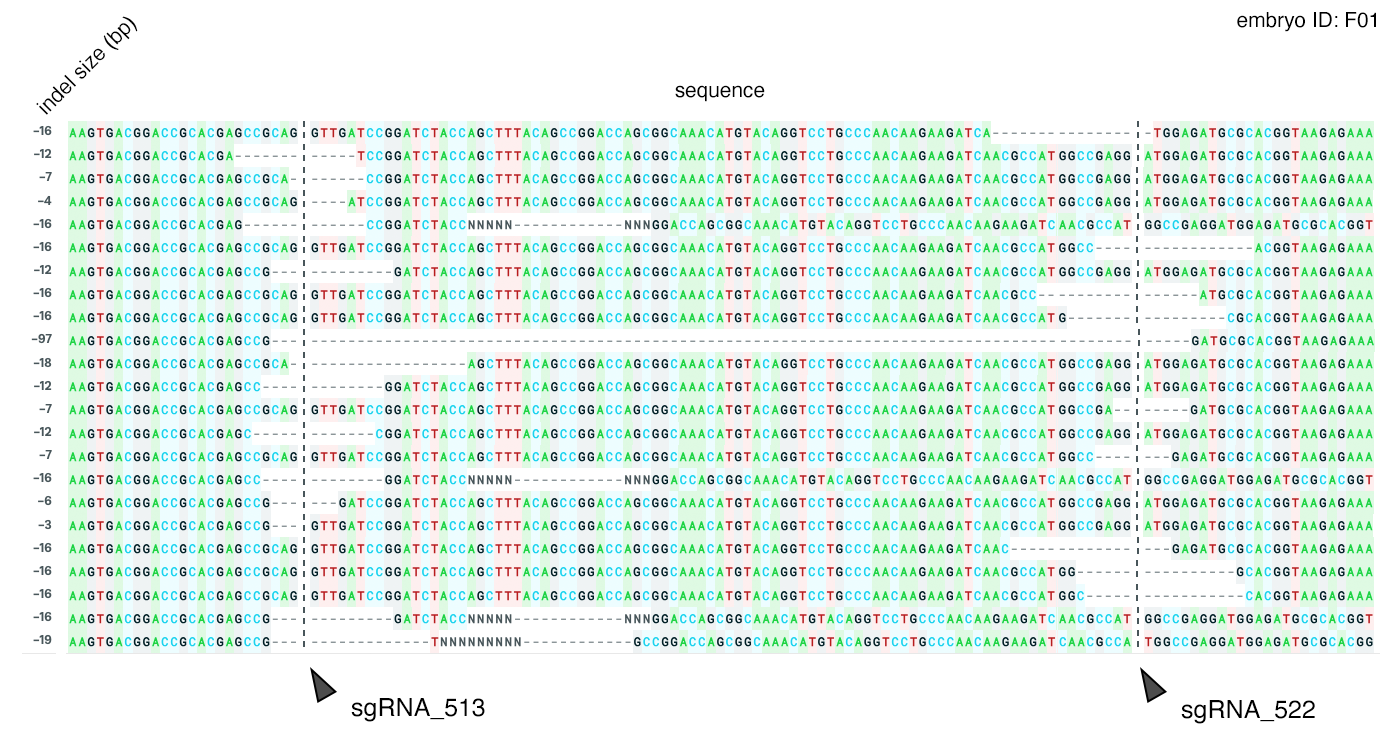

**Supplementary Figure 3:** Extended VISTA alignment of the *fgf8a* locus as presented in Figure 4 with additional teleost species.

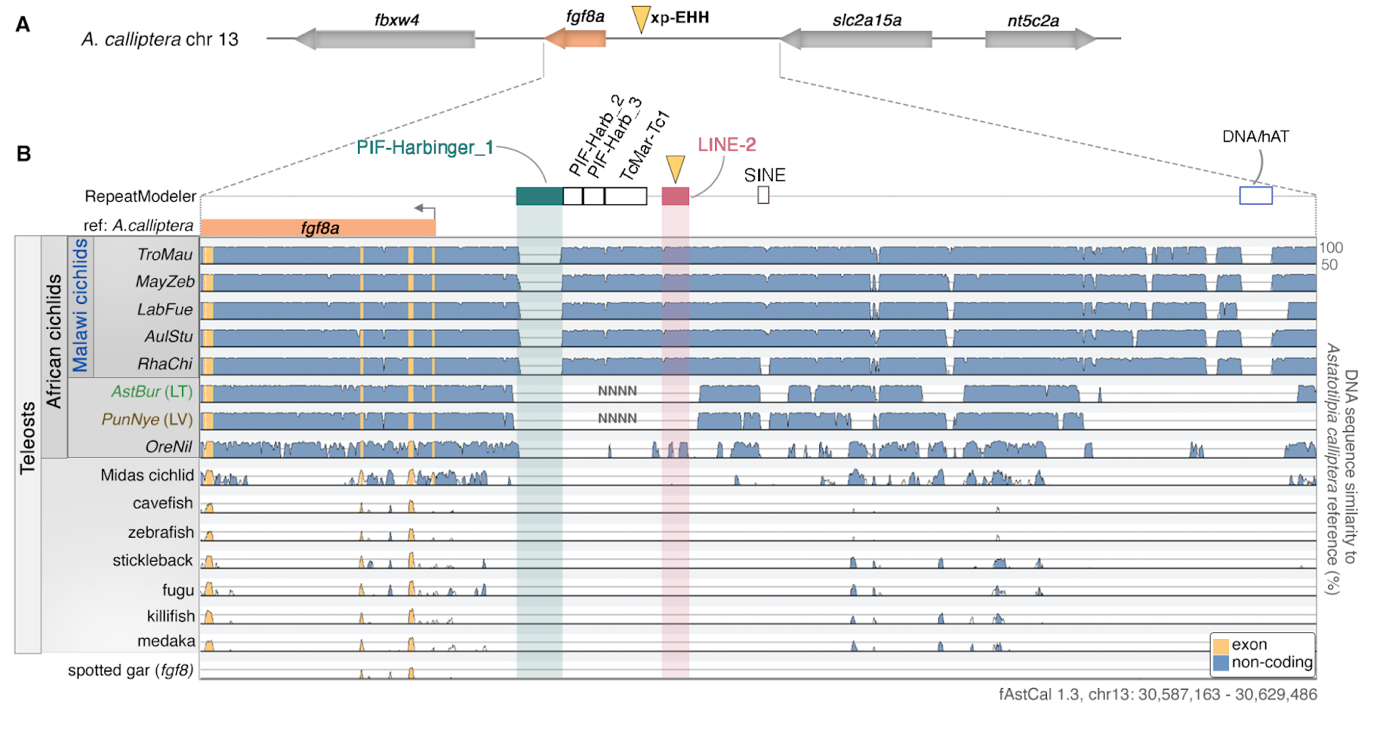

**Supplementary Figure 4:** Elevated rates of transposable element polymorphism in *fgf8a* locus in *Rhamphochromis* clade compared to the rest of the Lake Malawi radiation. A. Neighbour-Joining phylogeny of the Lake Malawi radiation based on the TE polymorphism within the 120kb interval spanning *fgf8a* and *slc2a15a*. B. The proportion of private TE polymorphisms in the *fgf8a* locus in *Rhamphochromis* is elevated compared to the radiation-wide average in equivalent windows on chromosome 13.

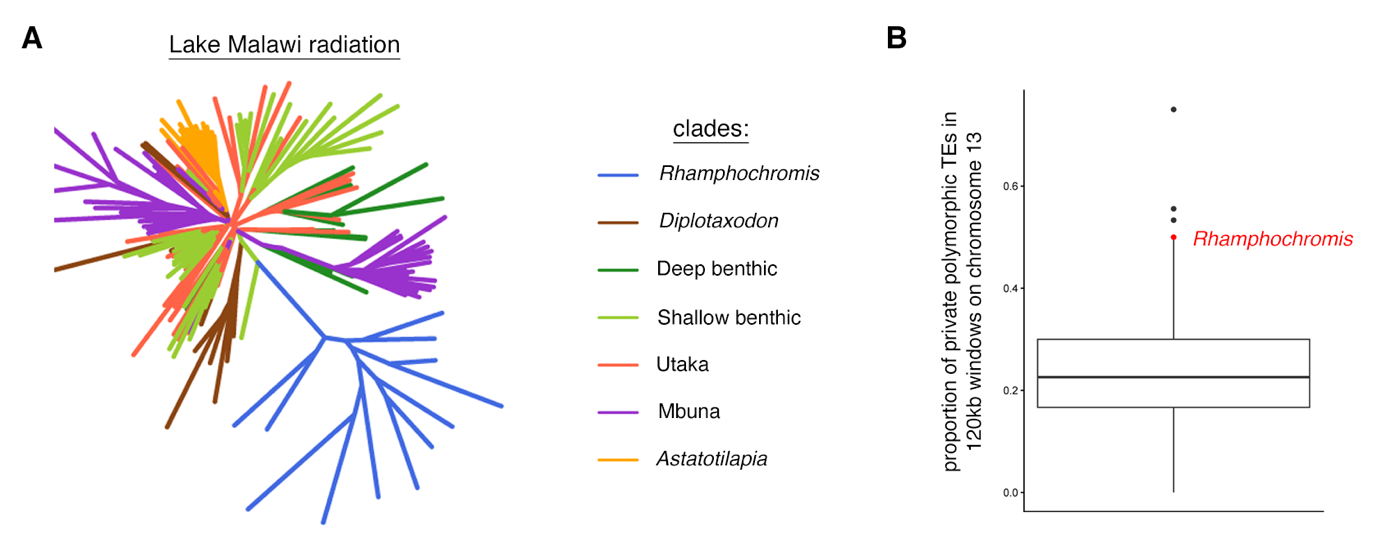

**Supplementary Figure 5:** Sequence alignment of LINE-2 transposable element insertion in *fgf8a* locus between *Astatotilapia calliptera* and *Rhamphochromis* sp. ‘chillingali’ highlights the 3 bp indel overlapping with a bHLH transcription factor binding site.

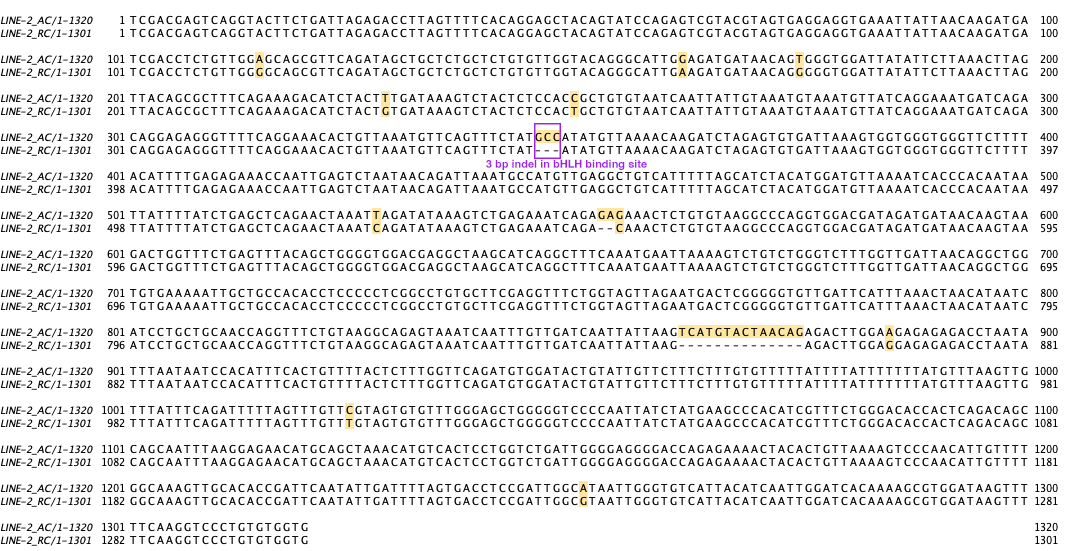

**Supplementary Tables List (supplied in a single Excel workbook)**

**Supplementary Table 1:** Summary of the CRISPR knockout experiments including survival rates and knockout efficiency.

**Supplementary Table 2:** Genomic coordinates of transposable element insertions within *fgf8a* locus from curated library.

**Supplementary Table 3:** Transcription factor binding site analysis of TE insertions in *fgf8a* locus using FIMO. Only significant results shown.

**Supplementary Table 4:** Summary of the zebrafish enhancer (reporter gene) assay.

**Supplementary Table 5:** Probe lot numbers for HCR probes purchased from Molecular Instruments.

**Supplementary Table 6:** Zebrafish gene identifiers used for Gene Ontology analysis of transcription factor binding sites found in TE sequences.

**Supplementary Data Files**

**Supplementary Data Files 1-3:** GenBank files containing plasmid sequences (enhancer reporter constructs) from whole-plasmid next-generation sequencing.
